# Galápagos yellow warblers differ in behavioural plasticity in response to traffic noise depending on proximity to road

**DOI:** 10.1101/2024.06.10.598213

**Authors:** Leon Hohl, Alper Yelimlieş, Çağlar Akçay, Sonia Kleindorfer

## Abstract

Acoustic communication between animals is increasingly disrupted by noise in human-altered environments making signals less effective. Birdsong is a signal used in agonistic interactions between territorial rivals, and birds may modify their song and singing behaviour in response to noise. However, if these modifications are still ineffective, this can lead to increased conflict between rivals. Here, we asked whether experimental traffic noise induces immediate changes in acoustic characteristics of song and aggressive behaviour in populations and territories that differ greatly in traffic noise exposure. We conducted simulated territorial intrusions on Galápagos yellow warblers (*Setophaga petechia aureola*) living on Santa Cruz (high traffic) and Floreana (low traffic) islands. Territories were either adjacent to the nearest road or at least 100 m away from it. We assessed the focal birds’ physical response levels and recorded their vocalisations in response to playback of conspecific song (control) and conspecific song coupled with traffic noise (noise treatment). We found that physical response levels mostly depended on treatment and distance to the nearest road: on both islands, birds living adjacent to the nearest road increased their aggression levels with experimental noise compared to control, while birds living farther away decreased their aggression levels. Birds on all islands irrespective of their distance to a road increased minimum frequency of their songs during the noise treatment. However, change in peak frequency and duration of their songs depended on the habitat they live in. Our results suggest behavioural flexibility in territorial responses and birdsong in response to traffic noise, which appears to depend at least in part on prior experience with traffic noise.

**Highlights:**

- Galápagos yellow warblers modified their behaviour in response to traffic noise.
- Behavioural plasticity depended on the island and distance to the nearest road.
- Birds living close to a road increased their aggression in response to an intruder
- Birds living farther away from a road decreased their aggression.

---

Interference from noise during signal transmission has important implications for animal communication (Bradbury & Vehrencamp, 2011). Given the rapidly increasing scale of human activities, animals need to produce and perceive signals in an ever-noisier world (Brumm, 2013; Halfwerk & Slabbekoorn, 2015). The need to cope with noise is especially prevalent in the acoustic modality, where loud noises caused by traffic, construction, or other activities may mask animal vocalisations with implications for lower reproductive success and/or survival (Bowen et al., 2020; De Jong et al., 2018; Grade & Sieving, 2016).

The effect of anthropogenic noise on vocalisations has been most widely studied in songbirds (Brumm & Zollinger, 2013; Duquette et al., 2021). Bird song is a culturally transmitted acoustic signal that functions in both inter-sexual communication such as mate attraction as well as communication within each sex, such as territory defence against same-sex competitors (Catchpole & Slater, 2008). Signaling in territorial disputes can be mutually beneficial, allowing both parties to assess their rivals without the cost of fighting, because fights may result in injury or death (Logue et al., 2010; Maynard Smith & Harper, 1988). When anthropogenic noise masks animal signals, however, assessment of rivals’ quality or motivation via signals is impaired, increasing the probability that the opponent will need to resort to a conflict involving physical aggression. In support of this scenario, there is evidence from several species of songbirds that high levels of acoustic noise are correlated with increased physical aggression (Akçay et al., 2020; Davies & Sewall, 2016; Phillips & Derryberry, 2018). Of course, increased aggressiveness in urban environments could also be explained by other variables such as resource availability (Foltz et al., 2015), but experimental studies have shown that at least in some cases noise can lead to increased territorial aggression in birds (Grabarczyk & Gill, 2019; Önsal et al., 2022; but see Kleist et al., 2016).

To improve transmission of signals in noise, songbirds may use one of several adjustments to their signals and signaling behaviours (Brumm & Zollinger, 2013; Kunc & Schmidt, 2021). One common adjustment is to increase the amplitude of the signal, known as the Lombard effect (Brumm & Zollinger, 2011; Hage et al., 2013; Kunc et al., 2022). Alternatively, they can increase the redundancy of the signal through repetition or by singing for longer durations (Brumm & Slater, 2006; Ríos-Chelén et al., 2013). Increasing redundancy would be potentially more effective in response to intermittent noise (as opposed to constant noise) as there would be a possibility of at least partially avoiding masking during signal production.

Because anthropogenic noise generally occupies lower frequencies, another possibility to reduce the effects of masking is to change the spectral parameters of the vocalisations (Roca et al., 2016; Slabbekoorn & Peet, 2003). One possibility is to raise the frequency of vocalisations so that there is less overlap with low-frequency noise. Such changes in frequency may come about through either ontogenic effects (Moseley et al., 2018; but see Liu et al., 2021), or through immediate and short-term plasticity (Bermúdez-Cuamatzin et al., 2011; Verzijden et al., 2010). It is worth noting that the changes in frequency may partly come about as a by-product of the Lombard effect as singing louder can lead to higher frequencies (Nemeth & Brumm, 2010).

Whether animals can cope with noise by changing their signaling may depend on previous experience with noise (Courter et al., 2020; Gallego-Abenza et al., 2020; LaZerte et al., 2016). There are only a few studies that have directly examined whether previous experience with noise affects short-term changes in song in response to acute noise. In one study, LaZerte and colleagues (2016) found that black-capped chickadees (*Poecile atricapillus*) shifted their song frequencies to higher frequencies in response to noise only if they already lived in noisier territories. By contrast, birds in quieter territories shifted their song frequencies downward. In a similar vein, urban, but not rural, white-crowned sparrows (*Zonotrichia leucophyrs*) produced songs with lower maximum frequencies and narrower bandwidths when exposed to experimental noise (Gentry et al., 2017). Given that urban birds live in noisier territories this finding again suggests a role for experience. In contrast to these results, other studies found that in at least some species, individuals living in rural or quieter areas can also respond to experimental noise with immediate plasticity in vocalisations (Bermúdez-Cuamatzin et al., 2011; Potvin & Mulder, 2013; Verzijden et al., 2010).

While long considered a natural laboratory to study natural selection (B. R. Grant & Grant, 1979; P. R. Grant & Grant, 2014), the Galápagos Archipelago is not spared from the human footprint and is now increasingly also a source of research into urban ecological frameworks that explore human-wildlife conflict and the evolutionary dynamics of endemic island systems. In Darwin’s finches, there is a growing body of research into the effects of urbanisation on altered foraging behaviour (De León et al., 2019; Rivkin et al., 2021), nesting success (Harvey et al., 2021), boldness (Gotanda, 2020), song phenotypes and aggressive behaviour (Colombelli-Négrel et al., 2023). Given the large growth in the human population on the Galápagos Islands (Kricher & Loughlin, 2022; INEC 2016), there is a concomitant increase in the number of trucks and cars, and as a result higher levels of noise as well as more roadkill (García-Carrasco et al., 2020). For example, between 1980 and 2013, there was a 57-fold increase in traffic on Santa Cruz Island (23 vehicles in 1980 and 1326 vehicles in 2013; Márquez 2000; García-Carrasco et al. 2020). The endemic Galápagos yellow warbler (*Setophaga petechia aureola*) is widely distributed across the archipelago, both on inhabited and uninhabited islands, and is the most commonly killed bird (> 70%) among roadkill on the heavily trafficked main road of Santa Cruz Island (García-Carrasco et al., 2020). Being commonly found across the human disturbance gradient, the Galápagos yellow warbler has the potential to be an important study system for investigating the impacts of human activities on island birds.

Here we test whether immediate plasticity in the face of experimental noise is dependent on previous experience with chronic noise in two island populations of the male Galápagos yellow warblers. More specifically, we aim to measure behavioural and song structure plasticity in response to a simulated intruder under traffic noise and to compare the magnitude of response between two islands (Santa Cruz and Floreana) that differ in human population size and traffic. We also examined whether birds in territories adjacent to the road will respond differently to intruders compared to birds in territories farther away from the nearest road. Roadside territories will be noisier and birds living in these territories will have direct previous experience with traffic noise. If the birds show behavioural and song structure plasticity in their response to an intruder song in a noisy environment, we expect that individuals will modify their responses. In particular they are expected to change their songs to increase detectability in noise by increasing frequency, duration and singing more songs. Crucially, if the ability to modify song structure depends on experience with noise (Gentry et al., 2017; LaZerte et al., 2016), more modifications will be made a) on the island with more traffic exposure (Santa Cruz) than less traffic exposure (Floreana), and b) in road-side territories compared to territories farther away from the road. Finally, we also expected that if birds do not modify their song or modifications are ineffective in dealing with experimentally increased noise, they will increase physical aggression by more closely approaching the simulated intruder and engaging in more search behaviour for the intruder. Again, if this effect depends on previous experience, we expect it to be more pronounced in the island with higher traffic and in roadside territories.

## Methods

### Study species and sites

The Galápagos yellow warbler is an abundant songbird that occurs on all vegetated islands and across all habitats of the Galápagos Archipelago, Ecuador (Chaves et al., 2012; Snow, 1966). It is a small (∼12 g) songbird that is territorial year-round (Snow, 1966). Both sexes sing, and the sexes are distinguishable by their songs (Yelimlies et al. unpublished data). Males have a reddish-brown crown and a chest streaking of the same colour against a yellow and olivaceous plumage, whereas females are only yellow and olivaceous (Salgado-Ortiz et al., 2008), making the sexes distinguishable using binoculars in the field.

We conducted the experiment in the lowlands and highlands on Santa Cruz Island and Floreana Island. All study sites were previously part of a long-term annual nesting survey of Darwin’s finch breeding biology that has been conducted by our research group since 2000 (Common et al., 2022; Kleindorfer et al., 2021). The study plots in the Santa Cruz highlands were located in the Los Gemelos area (−0.625982, −90.384829), which is away from most human settlements, but is crossed by the island’s main road with heavy traffic, and the vegetation is dominated by *Scalesia* forests. The lowland study plots were in the El Barranco area (0.739068, - 90.301467) near the buildings of Charles Darwin Foundation and are vegetated with cacti and shrubs. The Floreana Island highland study plots were located in the Cerro Pajas (−1.299829, - 90.455674) and Asilo de la Paz (−1.313595, −90.454935) areas. Both are dominated by *Scalesia* forests. Asilo de la Paz is a tortoise sanctuary with occasional tourist visits adjacent to agricultural land, whereas Cerro Pajas area is only accessible to the national park staff and researchers with permits. The lowland sites on Floreana island were near the Loberia area (−1.282974, −90.49208) near the only town of the island and mainly vegetated with *Opuntia* cacti and Palo Santo (*Bursera graveolens*) trees. Both islands have permanent human settlements but differ in human population size and number of vehicles (INEC 2016; Galápagos Conservancy 2023). Santa Cruz has a population of over 15,000 humans (INEC 2016) and over 1,000 vehicles (García-Carrasco et al. 2020) compared with Floreana which has a human population of ∼100 (Galápagos Conservancy 2023) and ∼10 vehicles (Hohl et al. personal observation). The current selection of the islands shows the highest traffic noise contrast possible on the inhabited islands but the Galápagos archipelago has two more inhabited islands (i.e. Isabela and San Cristobal), that could have been included in the present study. Unfortunately, due to logistical constraints, we were unable to do so, which represents a significant caveat.

### Number of vehicles

To estimate relative traffic density between the two islands, we placed passive sound recorders developed by the New Zealand Department of Conservation along the main road in the highlands at Los Gemelos on Santa Cruz (28 and 29 Jan) and Cerro Pajas on Floreana Island (4 and 5 Feb). The number of passing vehicles was counted by listening to the recordings at x8 speed on headphones of recordings made between 0600 and 0800 and 1000 and 1200. On 28 Jan on Santa Cruz Island, we measured max dB (A-weighted and fast response, 125 ms) in the field with a sound level meter (Voltcraft SL-100) standing roadside at 3 m from 27 passing vehicles which lasted for about ∼30 seconds.

### Active territories

We carried out playback trials at 38 (22 Floreana, 16 Santa Cruz) active yellow warbler territories in January and March 2022 on Santa Cruz, and February 2022 on Floreana. Focal birds were identified from 3-7 visits to the nest territories across three weeks. Sites were systematically searched every two days for the presence of male and female pairs and active nests (Lawson et al., 2023). A male or a male and female pair was considered resident if they were active on the territory over a three-day period as evidenced by nest building and/or males showing mate-guarding (following a female) and/or females exhibiting nest defence behaviours (alarm-calling, perch-switching, wing-flicking, circle-flight), which have been used to assign partulid nesting status in similar studies (Hobson & Sealy, 1989; Mitra, 1999). As in Lawson et al. (2023), if these conditions were met, the territory was included in the experiment

We took GPS measurements of each location where we carried out playback and classified them as either roadside (<50 m from the nearest road) or non-roadside (>100m from the nearest road, there were no territories situated between 50 and 100 meters away from the road in our sample). We had 10 roadside territories on each island and 6 and 12 non-roadside territories on Santa Cruz and Floreana islands, respectively; for a total of 38 territories.

### Playback stimuli

We prepared 15 song and 10 traffic noise stimulus files from our field recordings using the software Audacity. The 15 yellow warbler song stimuli (9 from Floreana, 6 from Santa Cruz) were created from recordings made in January and February 2022 using a Zoom H5 (Zoom Corporation, Japan) recorder with a Sennheiser MKE 600 (Sennheiser electronic GmbH & Co. KG, Germany) directional microphone. Recordings were made at a 48 kHz sampling rate and 24-bit resolution. We selected songs with a high signal-to-noise ratio from the recordings, filtered frequencies below 1500 Hz, and normalised the peak amplitude of each song. The song stimuli consisted of 2 or 3 renditions of a single song type presented at a rate of 6 songs per minute. This type of stimulus in which they repeat the same song type approximates their natural behaviour in agonistic interactions (i.e. Type 1 singing; Beebee, 2004).

We generated traffic noise stimulus files using recordings of vehicles passing on the main road through Santa Cruz Island during January 2022 by standing on the roadside and pointing the microphone towards the road (using the same equipment and settings as described above). We edited the traffic noise recordings to first clip unique traffic events (a vehicle passing). Then, we merged on average 5.5 (SD = 1.2) events per minute to create a stimulus tape of traffic noise. In other words, our noise stimulus simulated a traffic of 5.5 vehicles per minute passing by. This noise stimulus simulates the typical intermittent and varying levels of noise (which increases and decreases as a vehicle passes) that birds experience. Figure 1 shows the average frequency spectra for the noise and the song stimuli. Each playback stimulus consisted of 4 minutes: 1-minute baseline (silence), 1-minute song (song or song coupled with traffic noise, depending on treatment), 1-minute silence or traffic noise, and 1-minute song (song or song coupled with traffic noise). The song stimulus for each subject was a local song from a non-neighbouring conspecific recorded at least 200 m away from the same island. Both stimuli (song and noise) were broadcast at 85 dBA max (measured at 1 m from the speaker with Voltcraft SL-100 with the same settings as described above, Conrad Electronic, Germany).

**Figure 1.**
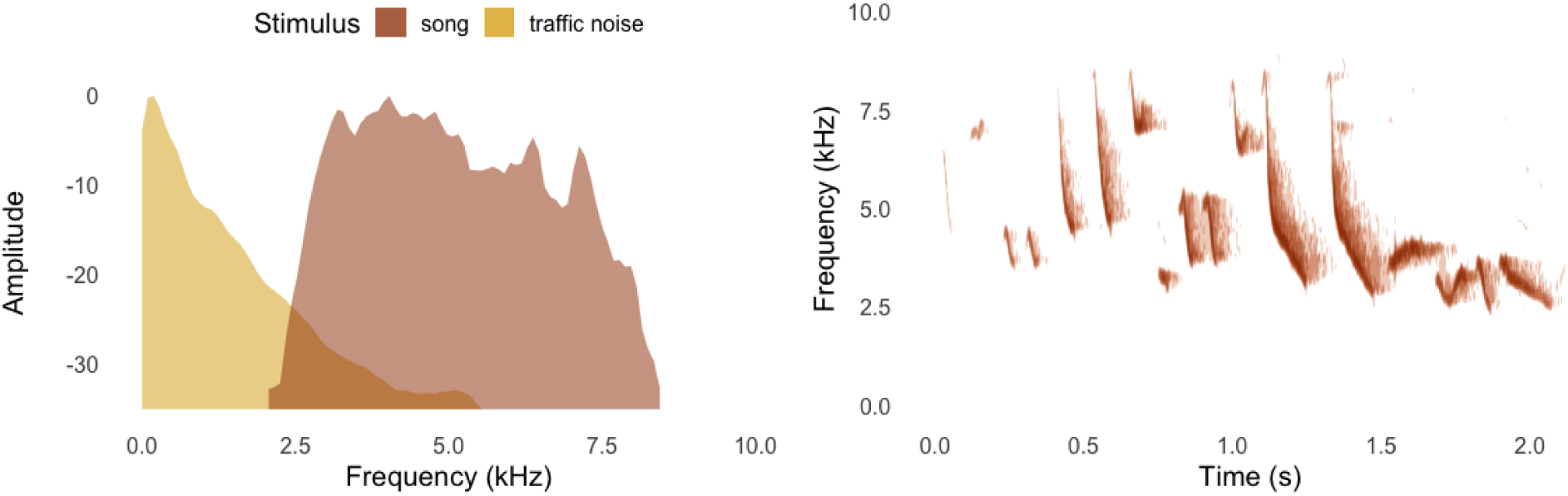
On the left: mean frequency spectra for one example traffic noise stimulus file and one song stimulus file, showing the overlap between the two. On the right: spectrogram of the same yellow warbler song stimulus.

### Playback procedure and response measures

Each male was tested twice, once with song only and once with song plus traffic noise, the two trials being 1 day apart except for the 3 birds that were tested 2 to 5 days apart due to rainy weather. 20 of the subjects were tested with the song-only treatment first and the remaining 18 were tested with the noise treatment first. For each trial, we placed two JBL Clip 4 Bluetooth speakers (JBL Inc., USA) at 0.5 m and 1.5 m above ground (usually clipped on a branch). The lower speaker was used to broadcast the traffic noise on the traffic noise trials and the upper one was used for the song. After placing the speakers, two observers were positioned 10 m apart and waited until the male was seen within 20 m to start the playback. Once the male was located within range, one observer started recording with a Zoom H5 recorder with a Sennheiser MKE 600 microphone and started the playback stimulus via a smartphone. During the trials, starting with the 1-minute baseline period, both observers narrated the movements of the focal bird and recorded songs. From the narrations, we extracted the closest approach distance, number of flights, and crosses to the speaker, defined as flights within a 1m radius of the conspecific playback speaker. These measures are commonly used to quantify physical aggressive responses in songbirds (Falls & Brooks, 1975; Searcy et al., 2006). We also counted the number of songs each male sang from these recordings.

### Song analysis

From the trial recordings, we extracted high-quality songs that did not overlap with other vocalisations using the software Raven Pro 1.6.4 (K. Lisa Yang Center for Conservation Bioacoustics, 2022) to quantify their song structure. The resulting dataset consisted of 490 songs (Santa Cruz = 225, Floreana = 265, 13.61 songs on average with a standard deviation of 8.65 per bird) from 36 birds (Santa Cruz = 15, Floreana = 21). We then measured the minimum and maximum frequency of the selected songs −20 dB relative to the peak frequency (i.e. the threshold method, Brumm et al., 2017); and calculated bandwidth as their difference using the ‘freq_detec’ function from the warbleR package in R (Araya-Salas & Vidaurre, 2017; R Core Team, 2023). We also measured the peak frequency and the duration of the songs within Raven Pro. For the latter, we used the duration 90% function which measures the duration in each selection that is between the times dividing the selection to the 5th and 95th percentile of the energy in the selection.

### Statistical analysis

All data were analysed using R version 4.3.2 (R Core Team, 2023). Because our measures of physical approach were all correlated with each other (minimum distance, crosses, flights) we used a principal component analysis (PCA) using the package psych in R (Revelle, 2017) for these variables. The first component (PC1) of the PCA (unrotated, using correlation matrix) explained 76% of the variance and had high factor loadings for all the behavioural responses associated with physical aggression as listed above so it was taken as the aggression scores (see Table S1 for factor loadings). A high aggression score (PC1) indicates a close approach and many flights and crosses over the speaker.

To test whether behavioural response and song structure differs when exposed to an intruder in a noisy environment depending on previous experience, we took a multi model inference approach. We first constructed linear mixed models for each response variable (aggression score, number of songs, and for the song structure: minimum frequency, maximum frequency, bandwidth, peak frequency, and duration) using the function ‘lmer’ from the package lme4 (Bates et al., 2015). The full models included treatment, island, territory location (roadside vs. non-roadside), all the two- and three-way interactions between these three variables and the treatment order as fixed effects, also bird ID as the random effect. We then averaged all the top models using ‘dredge’ and ‘model.avg’ functions from the MuMIn package (Bartoń, 2024). We took the full average of the models with a delta AICc’s < 6 following Harrison and colleagues (2018). For each variable present in the averaged model we reported averaged estimates, unconditional standard errors, 95% confidence intervals (95% CI) and relative variable importance (RVI, i.e. for each term, the sum of Akaike’s weights from the averaged models in which it appeared). We evaluated RVI scores and 95% confidence intervals to infer about the importance of a factor.

### Ethical note

This study complies with all current Austrian laws and regulations and was supported by Animal Experiment License Number 66.006/0026-WF/V/3b/2014 issued by the Austrian Federal Ministry for Science and Research (EU Standard, equivalent to the Animal Ethics Board). The subjects were not captured during the experiment and each trial only took four minutes. The birds returned to their normal activities within a few minutes.

## Results

### Traffic conditions between islands

There was significantly more vehicle traffic on Santa Cruz Island (62 ± 7.8 per hr) than on Floreana Island (1.8 ± 0.3 per hr, t = 7.679, df = 6, P < 0.001). The average amplitude of 27 vehicles recorded at 3 m was 84.7 ± 0.7 dBA max, with a minimum of 79.9 and a maximum of 95.4.

### Physical responses to playback

All yellow warblers approached the speaker and displayed territorial defence behaviours during the playbacks (Table S2). In our averaged model, the interaction between territory location and treatment was the most important effect explaining the aggression scores; yellow warblers increased their aggression levels during noise trials if their territory was on the roadside (<50m), but they decreased their aggression levels if their territory was far from the road (>100m). The interaction between island and treatment was also important, albeit the effect was weaker (see Table 1, Figure 2). Also, the treatment order influenced aggression scores, all birds being less aggressive during their second trials (Table 1).

**Table 1.** Averaged LMM for aggression score based on all the top models with ΔAICc< 6. All models included bird ID as the random factor. Variables for which the 95% confidence intervals of the coefficient does not include zero are bolded. RVI= relative variable importance.

| Fixed effects | Estimate | SE | 95% CI | RVI |
| --- | --- | --- | --- | --- |
| **Intercept** | 0.31 | 0.25 | [-0.17, 0.79] |  |
| **Island [Santa Cruz]** | 0.14 | 0.36 | [-0.57, 0.85] | 0.92 |
| **Territory location [roadside]** | -0.30 | 0.34 | [-0.96, 0.37] | 1.00 |
| **Treatment [Song+Noise]** | -0.30 | 0.20 | [-0.68, 0.09] | 1.00 |
| **Treatment order [second trial]** | **-0.51** | **0.12** | **[-0.73, -0.28]** | **1.00** |
| **Island [Santa Cruz] x treatment [Song+Noise]** | **-0.67** | **0.32** | **[-1.29, -0.06]** | **0.92** |
| **Territory location [roadside] x treatment [Song+Noise]** | **1.08** | **0.25** | **[0.6, 1.57]** | **1.00** |
| **Island [Santa Cruz] x territory location [roadside]** | 0.15 | 0.39 | [-0.63, 0.92] | 0.34 |
| **Island [Santa Cruz] x territory location [roadside] x treatment [Song+Noise]** | 0.05 | 0.21 | [-0.35, 0.46] | 0.11 |

**Figure 2.**
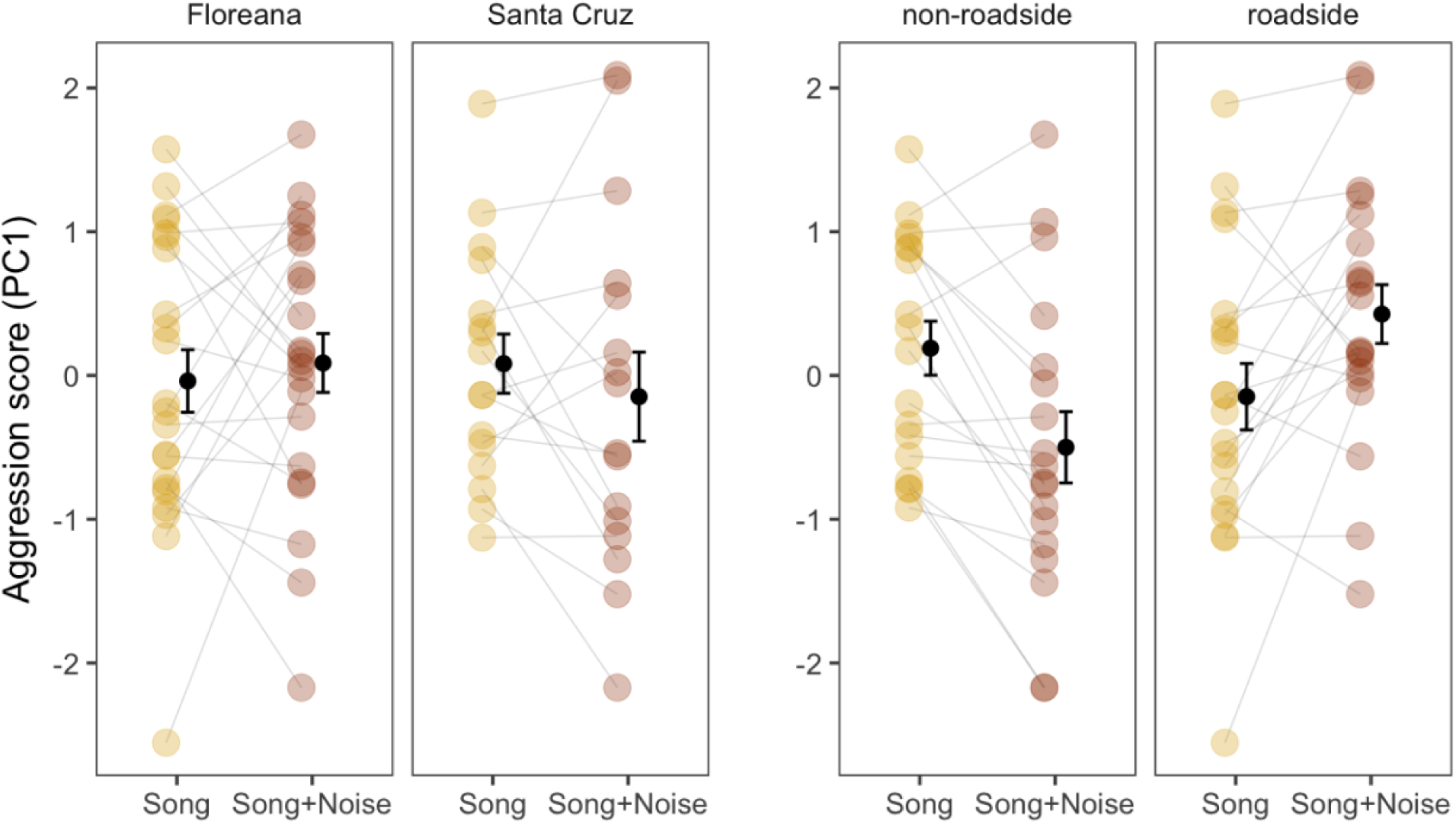
Change in response to treatment, showing mean ± SE for aggression scores (PC1) in control and experimental trials depending on island and distance to the road.

### Singing rate and song features

For the number of songs, none of the factors were important in our models, it did not differ by island, treatment, territory location, any interaction between them, or the treatment order (see Table S3). Males sang with slightly higher minimum frequencies in the second trials and during the noise treatment, but no other effects were important in explaining the variance in minimum frequency (see Table2a, Figure 3). The interaction between treatment and territory location was important for the models of peak frequency, males sang with a higher peak frequency during the noise treatment only if their territory was away (>100m) from the roadside (Table 2b, Figure 3). For song duration, the two-way interaction between treatment and island was important: birds on Santa Cruz increased the song duration in response experimental noise while birds on Floreana decreased their song duration (Table 2c, Figure 3). None of the factors were important in the models for maximum frequency and bandwidth (Tables S4 and S5).

**Table 2.** Outputs of the model-averaged LMM’s for song parameters based on all the top models with ΔAICc< 6. All models included bird ID as the random factor. Variables for which the 95% confidence intervals of the coefficient does not include zero are bolded. RVI= relative variable importance.

| **a) Minimum Frequency** |  |  |  |  |
| --- | --- | --- | --- | --- |
| **Fixed effects** | **Estimate** | **SE** | **95% CI** | **RVI** |
| **Intercept** | 2453.87 | 41.94 | [2371.66, 2536.08] |  |
| **Island [Santa Cruz]** | 96.84 | 71.02 | [-42.36, 236.03] | 0.84 |
| **Treatment [Song+Noise]** | **75.45** | **33.25** | **[10.28, 140.63]** | **1.00** |
| **Treatment order [second trial]** | **57.67** | **19.88** | **[18.7, 96.64]** | **1.00** |
| **Island [Santa Cruz] x treatment [Song+Noise]** | -46.85 | 52.90 | [-150.52, 56.83] | 0.59 |
| **Territory location [roadside]** | 5.59 | 40.38 | [-73.56, 84.74] | 0.44 |
| **Territory location [roadside] x treatment [Song+Noise]** | -8.92 | 28.96 | [-65.69, 47.85] | 0.17 |
| **Island [Santa Cruz] x territory location [roadside]** | -6.61 | 43.56 | [-91.98, 78.76] | 0.12 |
| **Island [Santa Cruz] x territory location [roadside] x treatment [Song+Noise]** | 6.51 | 35.17 | [-62.42, 75.44] | 0.04 |
| **b) Peak Frequency** |  |  |  |  |
| **Intercept** | 4384.98 | 103.66 | [4181.8, 4588.15] |  |
| **Territory location [roadside]** | 174.74 | 140.99 | [-101.6, 451.08] | 0.98 |
| **Treatment [Song+Noise]** | **418.25** | **115.36** | **[192.14, 644.36]** | **1.00** |
| **Territory location [roadside] x treatment [Song+Noise]** | **-433.73** | **153.85** | **[-735.26, -132.19]** | **0.98** |
| **Treatment order [second trial]** | 47.81 | 69.80 | [-89, 184.61] | 0.49 |
| **Island [Santa Cruz]** | 19.62 | 102.14 | [-180.57, 219.82] | 0.44 |
| **Island [Santa Cruz] x treatment [Song+Noise]** | 16.79 | 66.58 | [-113.7, 147.28] | 0.15 |
| **Island [Santa Cruz] x territory location [roadside]** | -10.17 | 89.49 | [-185.57, 165.23] | 0.12 |
| **c) Duration** |  |  |  |  |
| **Intercept** | 0.95 | 0.04 | [0.87, 1.04] |  |
| **Island [Santa Cruz]** | -0.07 | 0.07 | [-0.21, 0.07] | 1.00 |
| **Treatment [Song+Noise]** | **-0.09** | **0.03** | **[-0.16, -0.03]** | **1.00** |
| **Island [Santa Cruz] x treatment [Song+Noise]** | **0.16** | **0.05** | **[0.06, 0.25]** | **1.00** |
| **Treatment order [second trial]** | 0.00 | 0.01 | [-0.03, 0.02] | 0.30 |
| **Territory location [roadside]** | 0.00 | 0.05 | [-0.1, 0.09] | 0.49 |
| **Territory location [roadside] x treatment [Song+Noise]** | 0.01 | 0.03 | [-0.05, 0.07] | 0.22 |
| **Island [Santa Cruz] x territory location [roadside]** | -0.02 | 0.06 | [-0.14, 0.11] | 0.18 |
| **Island [Santa Cruz] x territory location [roadside] x treatment [Song+Noise]** | 0.00 | 0.02 | [-0.04, 0.05] | 0.03 |

**Figure 3.**
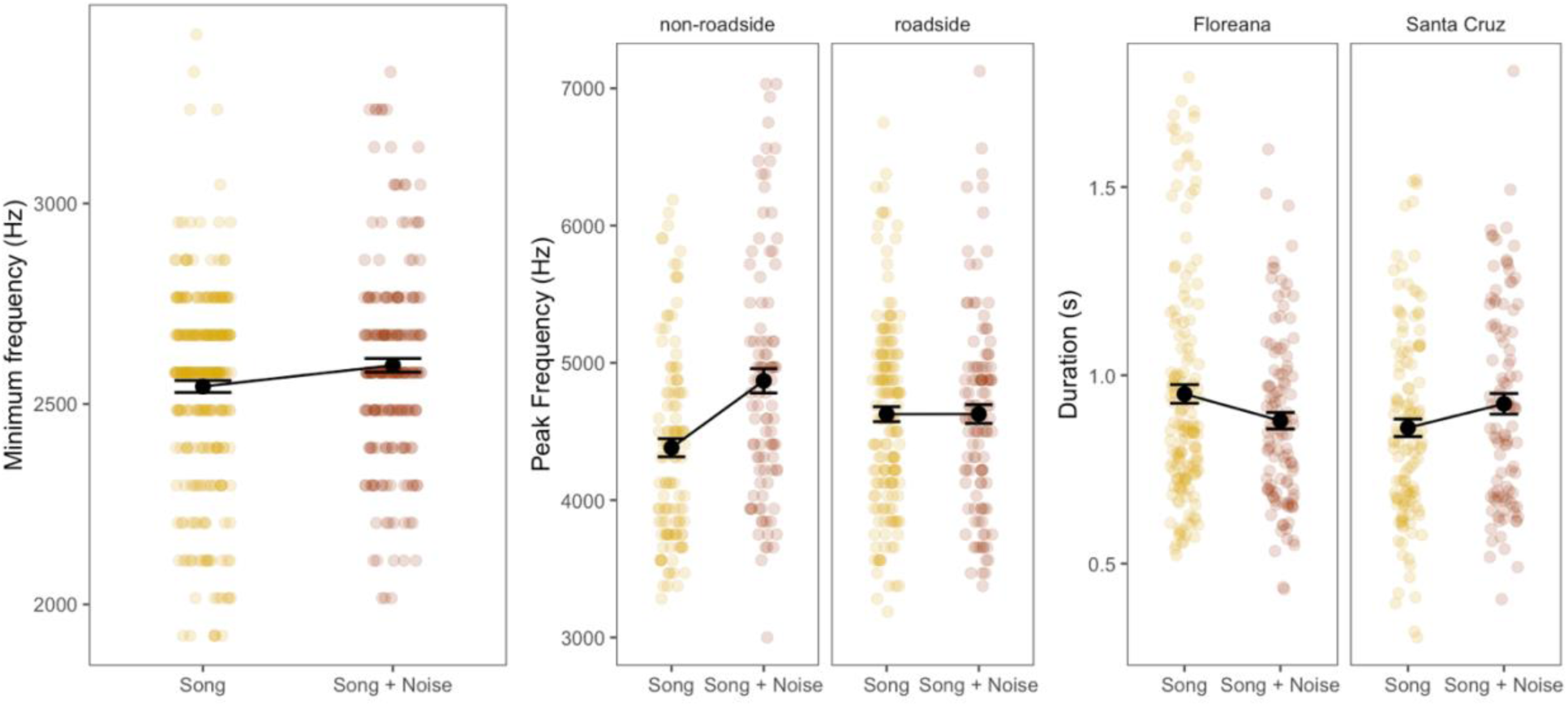
Changes in song structure variables. Each coloured circle represents a song, black dots in the middle and intervals show group means ± SE. Minimum frequency increased during the noise treatment, peak frequency increased during the noise treatment only in the non-roadside territories, and duration increased during the noise treatment only on the Santa Cruz (high traffic) island but decreased on the Floreana (low traffic) island.

## Discussion

In the present study, we asked whether Galápagos yellow warblers show increased aggression and flexibility in their song when confronted with a simulated intruder under conditions of experimentally increased noise depending on their experience with traffic noise. Our results showed that the change in aggressive responses in yellow warblers occurred mainly in proximity to the road: birds occupying roadside territories on both islands (and therefore presumably having regular experience of traffic noise on their territories) increased physical aggression in response to a simulated intrusion with experimental noise compared to without experimental noise. In contrast, birds occupying territories at least 100 m away from the nearest road decreased their aggression levels in response to intrusions with experimental noise compared to those without experimental noise. Surprisingly, the effect of island did not interact with this effect of proximity to the road, despite a large difference in traffic between the islands. We also found a slight increase in the minimum frequencies during the noise treatment irrespective of island and territory location. However, peak frequency of songs increased with experimental noise only in territories away from the road. Similarly, the song duration increased with experimental noise compared to the control treatment on Santa Cruz island, whereas song duration decreased in response to noise on Floreana island. The other acoustic parameters (maximum frequency and bandwidth) did not differ between the islands, proximity to road, or treatment. We discuss these results below.

If song produced during territorial defence is an aggressive signal that is used to resolve agonistic interactions instead of physical aggression, then acoustic noise that interferes with effective signaling may result in increased aggression. Several studies across taxa have found increased levels of aggression in noisy environments (in humans: Alimohammadi et al., 2018; Moudon, 2009; in songbirds: Akçay et al., 2020; Phillips & Derryberry, 2018). Here we found that experimentally presented traffic noise led to an increase in aggression levels only for the roadside territories but a decrease in territories away from the road. There was also a slight change in the opposite direction between islands in response to immediate noise: birds in the high-traffic island decreased aggression while they increased it in the low-traffic island. Nevertheless, this island and treatment interaction effect was much smaller and less robust then the first interaction between territory location and treatment. It is also worth noting that because we only tested two islands with differing levels of traffic, the island effect should be treated with additional caution. Future studies testing birds on additional islands differing in traffic levels would be needed to reliably examine the effect of traffic experience. We therefore conclude that, given the strong effect of proximity to road on both islands, even the slightest experience in traffic may matter for plasticity in aggressive behaviour.

Other studies testing the effect of immediate experimental noise on aggressive behaviour in birds yielded mixed results with an increase (Grabarczyk & Gill, 2019; Kort et al., 2024), no change (Akçay & Beecher, 2019; Zwart et al., 2016) or a decrease (Kleist et al. 2016; Reed et al., 2021). One other study on European robins (*Erithacus rubecula*) investigated whether this effect depends on the previous experience with noise (urban vs. rural living birds). Önsal and colleagues (2022) found that European robins without much anthropogenic noise experience (rural territories) rather than experienced ones (urban territories) increased aggression in response to immediate experimental noise. This difference in results between studies might be due to species and/or design differences. More specifically, Önsal and colleagues (2022) used a continuous noise filtered with the frequency spectrum of the city noise whereas in the present study we used fluctuating noise of cars passing by. A continuous low frequency noise might be more familiar to birds living away from urban noise because of its similarity to wind or river noise, however a burst of car noise might have caused a neophilia response to non-roadside birds in our study. For the urban birds, they argue that aggression levels being already higher without the experimental noise could be the reason why the urban European robins didn’t increase aggression. However, this baseline difference in aggression was not present with respect to the distance to road in our study, hence roadside birds increasing aggressive behaviours might be an appropriate response given the communication channel is blocked by noise.

We also found some evidence in favour of birds trying to cope with noise by adjusting song parameters. Yellow warblers in all habitats and on both islands slightly increased the minimum frequency of their songs in response to the experimental broadcast of noise, which was shown in other songbird species as well (Bermúdez-Cuamatzin et al., 2011; Verzijden et al., 2010). Birds on Santa Cruz increased the duration of their song more than the birds on Floreana, consistent with the hypothesis that long-term selection due to, or previous individual experience with noise, leads to the ability to rapidly and adaptively adjust songs features (Gentry et al., 2017; LaZerte et al., 2016). On the other hand, the peak frequency patterns that show an increase in non-roadside territories than in roadside ones appear to run against this hypothesis. A potential reason behind these seemingly contradictory findings may be that there is a limit to the benefit of increasing peak frequency; perhaps birds in roadside territories are already singing at the optimum peak frequency and further increasing it would result in a loss in the active space of the signal as high frequencies attenuate faster.

It is worth noting that the changes in frequency, while potentially adaptive, may also come about by increases in song amplitude. Nemeth and colleagues (2013) showed that amplitude and frequency covary and that birds with greater song amplitude also have higher frequencies. In our study, we did not measure the amplitude of the songs, and therefore can’t rule out an effect of the amplitude changes on song frequencies. Nemeth and Brumm, (2010) found that increasing amplitude is more effective in overcoming masking effects in noisy environments, which they measured experimentally in the great tit (*Parus major*) and blackbird (*Turdus merula*) populations. Specifically, higher amplitude songs travelled further whereas there was greater attenuation of higher pitched songs. In regard to the argument of efficacy, in urban habitats, the upward shift in song frequency increased the communication distance to a significantly lesser degree than amplitude (Nemeth & Brumm, 2010). Measuring amplitude in the field with freely moving animals is difficult but future studies should endeavour to disentangle these effects. In any case, although we found some seemingly adaptive responses to noise in song parameters, we do not know if these actually restore communication. So, another fruitful avenue would be to measure the transmission of the songs from the two islands under different noise treatments to see if the frequency as well as duration adjustments do in fact lead to better transmission.

In summary, we found evidence that noise-dependent plasticity in both aggressive behaviours and song parameters likely depends on the habitat experienced by animals. The results open new avenues of research questions regarding what aspects of experience shape behavioural plasticity in human-altered environments. In addition, while the Galápagos Islands are a natural living laboratory that inspired Charles Darwin’s formulation of the theory of evolution by natural selection (Darwin, 1859; P. R. Grant & Grant, 2014), our study adds to the growing body of knowledge that the archipelago is affected by anthropogenic impacts that are expanding globally. The Galápagos yellow warbler, common in all the habitats of vegetated islands, is a good model system for studying the human impacts on this unique archipelago.

## Data availability

Data and R-code to reproduce analyses can be found in the supplementary materials.

