## Supplementary file for "Galápagos yellow warblers differ in behavioural plasticity in response to traffic noise depending on proximity to road"

**Supplementary Materials**

**Table S1.** Factor loadings for the aggression scores (PC1)

|  | **Loading coefficients** |
| --- | --- |
| **Closest approach** | -0.77 |
| **#Crosses over speaker** | 0.91 |
| **#Flights** | 0.93 |

**Table S2.** The behaviour shown as number of songs and movement events per minute (mean ± SE) for Yellow Warbler males during a) the 1-minute period before the broadcast of the playback stimuli, and b) during broadcast of either the song or song + noise treatments. On both islands, resident males increased their singing behavior and movement in response to the broadcast of song of a simulated intruder with or without noise.

| **a)**  Pre-playback behaviour | | | | |
| --- | --- | --- | --- | --- |
|  | **Floreana** | | **Santa Cruz** | |
|  | **Song** | **Song + Noise** | **Song** | **Song + Noise** |
| **# Songs** | 8.6 ± 1.3 | 8.5 ± 0.9 | 11.1 ± 1.6 | 11.2 ± 1.6 |
| **#Crosses** | 3.9 ± 0.8 | 4.1 ± 0.7 | 4.1 ± 1.0 | 3.8 ± 1.2 |
| **#Flights** | 11.9 ± 1.8 | 12.9 ± 1.7 | 11.8 ± 2.0 | 11.4 ± 2.6 |
| **b)**  Playback behaviour | | | | |
|  | **Floreana** | | **Santa Cruz** | |
|  | **Song** | **Song + Noise** | **Song** | **Song + Noise** |
| **# Songs** | 8.6 ± 1.3 | 8.5 ± 0.9 | 11.1 ± 1.6 | 11.2 ± 1.6 |
| **#Crosses** | 3.9 ± 0.8 | 4.1 ± 0.7 | 4.1 ± 1.0 | 3.8 ± 1.2 |
| **#Flights** | 11.9 ± 1.8 | 12.9 ± 1.7 | 11.8 ± 2.0 | 11.4 ± 2.6 |

**Table S3.** Outputs of the model-averaged LMM for the number of songs produced during playbacks based on all the top models with ∆AICc< =6. All models included bird ID as the random factor.

| **Fixed effects** | **Estimate** | **SE** | **95% CI** | **RVI** |
| --- | --- | --- | --- | --- |
| (Intercept) | 7.88 | 1.34 | [5.26, 10.5] |  |
| islandSanta Cruz | 4.78 | 2.44 | [0, 9.56] | 0.92 |
| territory_locationroadside | -0.26 | 1.61 | [-3.43, 2.9] | 0.83 |
| treatment_ordersecond trial | 1.74 | 1.47 | [-1.15, 4.62] | 0.73 |
| islandSanta Cruz:territory_locationroadside | -3.59 | 3.28 | [-10.01, 2.83] | 0.65 |
| treatmentSong+Noise | -0.08 | 0.69 | [-1.44, 1.28] | 0.24 |
| territory_locationroadside:treatmentSong+Noise | 0.06 | 0.55 | [-1.02, 1.14] | 0.03 |
| islandSanta Cruz:treatmentSong+Noise | 0.03 | 0.44 | [-0.82, 0.88] | 0.03 |

**Table S4.** Outputs of the model-averaged LMM for the maximum frequencies of songs based on all the top models with ∆AICc< =6. All models included bird ID as the random factor.

| **Fixed effects** | **Estimate** | **SE** | **95% CI** | **RVI** |
| --- | --- | --- | --- | --- |
| (Intercept) | 7127.51 | 112.68 | [6906.66, 7348.36] |  |
| treatmentSong + Noise | 76.30 | 76.67 | [-73.97, 226.58] | 0.72 |
| treatment_ordersecond trial | 26.08 | 50.51 | [-72.92, 125.07] | 0.39 |
| territory_locationroadside | -24.38 | 124.07 | [-267.56, 218.81] | 0.41 |
| islandSanta Cruz | -6.89 | 138.31 | [-277.98, 264.19] | 0.40 |
| islandSanta Cruz:treatmentSong + Noise | 11.47 | 51.46 | [-89.39, 112.34] | 0.10 |
| territory_locationroadside:treatmentSong + Noise | -2.17 | 31.90 | [-64.7, 60.36] | 0.06 |
| islandSanta Cruz:territory_locationroadside | -42.36 | 163.54 | [-362.89, 278.18] | 0.09 |

**Table S5.** Outputs of the model-averaged LMM for the bandwidth of songs based on all the top models with ∆AICc< =6. All models included bird ID as the random factor.

| **Fixed effects** | **Estimate** | **SE** | **95% CI** | **RVI** |
| --- | --- | --- | --- | --- |
| Intercept) | 4646.02 | 117.22 | [4416.27, 4875.78] |  |
| islandSanta Cruz | -70.24 | 167.55 | [-398.62, 258.15] | 0.52 |
| treatmentSong + Noise | 8.82 | 56.91 | [-102.73, 120.37] | 0.48 |
| territory_locationroadside | -29.31 | 128.54 | [-281.24, 222.63] | 0.41 |
| islandSanta Cruz:treatmentSong + Noise | 34.40 | 88.66 | [-139.37, 208.17] | 0.18 |
| treatment_ordersecond trial | 1.75 | 29.64 | [-56.34, 59.84] | 0.25 |
| islandSanta Cruz:territory_locationroadside | -41.96 | 162.55 | [-360.55, 276.64] | 0.10 |
| territory_locationroadside:treatmentSong + Noise | 0.42 | 24.35 | [-47.29, 48.14] | 0.04 |

**Table S6.** Model averaging table for the LMM for aggression score as the output of dredge function in R. isl: island, loc: territory location, trt: treatment, ord: treatment order. Plusses indicate that the term was included in that model. Only the models with delta AICc <6 are shown in the table. Intercept and bird ID as the random effect was included in each model. ICC for the random effect was 0.72 in the top model.

| isl | loc | trt | ord | Isl:loc | Isl:trt | loc:trt | Isl:loc:trt | df | logLik | AICc | delta | weight |
| --- | --- | --- | --- | --- | --- | --- | --- | --- | --- | --- | --- | --- |
| + | + | + | + |  | + | + |  | 9 | -84.56805 | 189.8634 | 0.000000 | 0.5569121 |
| + | + | + | + | + | + | + |  | 10 | -84.14828 | 191.6812 | 1.817793 | 0.2244182 |
| + | + | + | + | + | + | + | + | 11 | -83.57123 | 193.2675 | 3.404087 | 0.1015310 |
|  | + | + | + |  |  | + |  | 7 | -89.09211 | 193.8313 | 3.967910 | 0.0765889 |

**Table S7.** Model averaging table for the LMM for number of songs produced during the playback as the output of dredge function in R. isl: island, loc: territory location, trt: treatment, ord: treatment order. Plusses indicate that the term was included in that model. Only the models with delta AICc <6 are shown in the table. Intercept and bird ID as the random effect was included in each model. ICC for the random effect was 0 in the top model as bird ID did not account for any variance.

| isl | loc | trt | ord | Isl:loc | Isl:trt | loc:trt | Isl:loc:trt | df | logLik | AICc | delta | weight |
| --- | --- | --- | --- | --- | --- | --- | --- | --- | --- | --- | --- | --- |
| + | + |  | + | + |  |  |  | 7 | -231.8795 | 479.4061 | 0.000000 | 0.26892327 |
| + | + |  |  | + |  |  |  | 6 | -233.8639 | 480.9452 | 1.539063 | 0.12457336 |
| + | + | + | + | + |  |  |  | 8 | -231.8681 | 481.8854 | 2.479300 | 0.07784939 |
| + | + |  | + |  |  |  |  | 6 | -234.3588 | 481.9350 | 2.528832 | 0.07594504 |
| + |  |  | + |  |  |  |  | 5 | -235.7046 | 482.2664 | 2.860240 | 0.06434802 |
| + | + |  |  |  |  |  |  | 5 | -236.2208 | 483.2988 | 3.892686 | 0.03840099 |
| + | + | + |  | + |  |  |  | 7 | -233.8630 | 483.3730 | 3.966821 | 0.03700362 |
| + |  |  |  |  |  |  |  | 4 | -237.5122 | 483.5877 | 4.181543 | 0.03323670 |
| + | + | + | + | + |  | + |  | 9 | -231.5866 | 483.9005 | 4.494400 | 0.02842378 |
|  |  |  | + |  |  |  |  | 4 | -237.7742 | 484.1117 | 4.705553 | 0.02557588 |
| + | + | + | + | + | + |  |  | 9 | -231.7706 | 484.2685 | 4.862370 | 0.02364711 |
| + | + | + | + |  |  |  |  | 7 | -234.3481 | 484.3432 | 4.937051 | 0.02278040 |
| + |  | + | + |  |  |  |  | 6 | -235.6940 | 484.6055 | 5.199323 | 0.01998066 |
|  | + |  | + |  |  |  |  | 5 | -237.0737 | 485.0045 | 5.598337 | 0.01636685 |

**Table S8.** Model averaging table for the LMM for the minimum frequencies of songs as the output of dredge function in R. isl: island, loc: territory location, trt: treatment, ord: treatment order. Plusses indicate that the term was included in that model. Only the models with delta AICc <6 are shown in the table. Intercept and bird ID as the random effect was included in each model. ICC for the random effect was 0.32 in the top model.

| isl | loc | trt | ord | Isl:loc | Isl:trt | loc:trt | Isl:loc:trt | df | logLik | AICc | delta | weight |
| --- | --- | --- | --- | --- | --- | --- | --- | --- | --- | --- | --- | --- |
| + |  | + | + |  | + |  |  | 7 | -3317.716 | 6649.663 | 0.000000 | 0.29388199 |
| + |  | + | + |  |  |  |  | 6 | -3319.535 | 6651.243 | 1.579652 | 0.13340001 |
| + | + | + | + |  | + |  |  | 8 | -3317.713 | 6651.725 | 2.061707 | 0.10482841 |
|  |  | + | + |  |  |  |  | 5 | -3321.000 | 6652.124 | 2.460655 | 0.08587141 |
| + | + | + | + |  | + | + |  | 9 | -3317.454 | 6653.283 | 3.619336 | 0.04811097 |
| + | + | + | + |  |  |  |  | 7 | -3319.535 | 6653.301 | 3.638068 | 0.04766246 |
| + | + | + | + | + | + |  |  | 9 | -3317.666 | 6653.706 | 4.043001 | 0.03892659 |
| + | + | + | + | + | + | + | + | 11 | -3315.587 | 6653.725 | 4.062120 | 0.03855625 |
|  | + | + | + |  |  |  |  | 6 | -3320.925 | 6654.025 | 4.361292 | 0.03319942 |
| + | + | + | + |  |  | + |  | 8 | -3318.924 | 6654.147 | 4.483475 | 0.03123192 |
|  | + | + | + |  |  | + |  | 7 | -3320.221 | 6654.674 | 5.010689 | 0.02399472 |
| + | + | + | + | + | + | + |  | 10 | -3317.418 | 6655.296 | 5.632656 | 0.01758155 |
| + | + | + | + | + |  |  |  | 8 | -3319.513 | 6655.325 | 5.661962 | 0.01732581 |

**Table S9.** Model averaging table for the LMM for the maximum frequencies of songs as the output of dredge function in R. isl: island, loc: territory location, trt: treatment, ord: treatment order. Plusses indicate that the term was included in that model. Only the models with delta AICc <6 are shown in the table. Intercept and bird ID as the random effect was included in each model. ICC for the random effect was 0.32 in the top model.

| isl | loc | trt | ord | Isl:loc | Isl:trt | loc:trt | Isl:loc:trt | df | logLik | AICc | delta | weight |
| --- | --- | --- | --- | --- | --- | --- | --- | --- | --- | --- | --- | --- |
|  |  | + |  |  |  |  |  | 4 | -3879.345 | 7766.773 | 0.000000 | 0.155356008 |
|  |  | + | + |  |  |  |  | 5 | -3878.887 | 7767.898 | 1.125594 | 0.088492918 |
|  | + | + |  |  |  |  |  | 5 | -3879.126 | 7768.375 | 1.602633 | 0.069714104 |
|  |  |  |  |  |  |  |  | 3 | -3881.233 | 7768.515 | 1.742262 | 0.065013082 |
| + |  | + |  |  |  |  |  | 5 | -3879.236 | 7768.596 | 1.823736 | 0.062417855 |
|  |  |  | + |  |  |  |  | 4 | -3880.357 | 7768.797 | 2.024368 | 0.056460150 |
|  | + | + | + |  |  |  |  | 6 | -3878.644 | 7769.463 | 2.689902 | 0.040478386 |
| + |  | + | + |  |  |  |  | 6 | -3878.772 | 7769.718 | 2.945794 | 0.035616969 |
| + |  | + |  |  | + |  |  | 6 | -3878.808 | 7769.789 | 3.016416 | 0.034381251 |
|  | + |  |  |  |  |  |  | 4 | -3880.964 | 7770.011 | 3.238087 | 0.030774178 |
|  | + |  | + |  |  |  |  | 5 | -3880.062 | 7770.248 | 3.474982 | 0.027336650 |
|  | + | + |  |  |  | + |  | 6 | -3879.065 | 7770.304 | 3.531576 | 0.026573944 |
| + | + | + |  |  |  |  |  | 6 | -3879.072 | 7770.319 | 3.545961 | 0.026383505 |
| + |  |  |  |  |  |  |  | 4 | -3881.136 | 7770.354 | 3.581437 | 0.025919629 |
| + | + | + |  | + |  |  |  | 7 | -3878.129 | 7770.490 | 3.717425 | 0.024215833 |
| + |  |  | + |  |  |  |  | 5 | -3880.251 | 7770.626 | 3.853627 | 0.022621614 |
| + |  | + | + |  | + |  |  | 7 | -3878.268 | 7770.768 | 3.995206 | 0.021075609 |
| + | + | + | + |  |  |  |  | 7 | -3878.589 | 7771.411 | 4.638096 | 0.015281930 |
| + | + | + | + | + |  |  |  | 8 | -3877.596 | 7771.491 | 4.718130 | 0.014682468 |
|  | + | + | + |  |  | + |  | 7 | -3878.639 | 7771.511 | 4.738129 | 0.014536386 |
| + | + | + |  |  | + |  |  | 7 | -3878.663 | 7771.559 | 4.786523 | 0.014188868 |
| + | + | + |  | + | + |  |  | 8 | -3877.788 | 7771.875 | 5.102041 | 0.012118086 |
| + | + |  |  | + |  |  |  | 6 | -3879.881 | 7771.937 | 5.163855 | 0.011749278 |
| + | + |  |  |  |  |  |  | 5 | -3880.924 | 7771.972 | 5.199054 | 0.011544307 |
| + | + |  | + | + |  |  |  | 7 | -3878.922 | 7772.077 | 5.303929 | 0.010954551 |
| + | + |  | + |  |  |  |  | 6 | -3880.018 | 7772.209 | 5.436715 | 0.010250862 |
| + | + | + |  |  |  | + |  | 7 | -3879.007 | 7772.246 | 5.472859 | 0.010067271 |
| + | + | + |  | + |  | + |  | 8 | -3878.081 | 7772.461 | 5.688650 | 0.009037605 |
| + | + | + | + |  | + |  |  | 8 | -3878.106 | 7772.511 | 5.738233 | 0.008816306 |
| + | + | + | + | + | + |  |  | 9 | -3877.185 | 7772.744 | 5.971708 | 0.007844912 |

**Table S10.** Model averaging table for the LMM for the bandwidth frequencies of songs as the output of dredge function in R. isl: island, loc: territory location, trt: treatment, ord: treatment order. Plusses indicate that the fixed factor or interaction term was included that model. Only the models with delta AICc <6 are shown in the table. Intercept and bird ID as the random effect was included in each model. ICC for the random effect was 0.35 in the top model.

| isl | loc | trt | ord | Isl:loc | Isl:trt | loc:trt | Isl:loc:trt | df | logLik | AICc | delta | weight |
| --- | --- | --- | --- | --- | --- | --- | --- | --- | --- | --- | --- | --- |
|  |  |  |  |  |  |  |  | 3 | -3864.974 | 7735.997 | 0.000000 | 0.142495187 |
| + |  |  |  |  |  |  |  | 4 | -3864.477 | 7737.037 | 1.040541 | 0.084693413 |
|  |  | + |  |  |  |  |  | 4 | -3864.564 | 7737.210 | 1.212700 | 0.077708009 |
|  | + |  |  |  |  |  |  | 4 | -3864.623 | 7737.329 | 1.331844 | 0.073213953 |
| + |  | + |  |  | + |  |  | 6 | -3862.667 | 7737.508 | 1.511505 | 0.066923870 |
|  |  |  | + |  |  |  |  | 4 | -3864.961 | 7738.004 | 2.006795 | 0.052243263 |
| + |  | + |  |  |  |  |  | 5 | -3864.052 | 7738.228 | 2.231677 | 0.046687185 |
|  | + | + |  |  |  |  |  | 5 | -3864.238 | 7738.600 | 2.603219 | 0.038772011 |
| + | + |  |  |  |  |  |  | 5 | -3864.276 | 7738.676 | 2.679269 | 0.037325389 |
| + | + |  |  | + |  |  |  | 6 | -3863.391 | 7738.956 | 2.959656 | 0.032442861 |
| + |  |  | + |  |  |  |  | 5 | -3864.462 | 7739.048 | 3.051199 | 0.030991370 |
|  |  | + | + |  |  |  |  | 5 | -3864.563 | 7739.251 | 3.253767 | 0.028006171 |
| + | + | + |  |  | + |  |  | 7 | -3862.523 | 7739.278 | 3.280781 | 0.027630435 |
|  | + |  | + |  |  |  |  | 5 | -3864.606 | 7739.337 | 3.339919 | 0.026825391 |
| + |  | + | + |  | + |  |  | 7 | -3862.660 | 7739.552 | 3.555634 | 0.024082645 |
| + | + | + |  | + | + |  |  | 8 | -3861.792 | 7739.884 | 3.886844 | 0.020407167 |
| + | + | + |  |  |  |  |  | 6 | -3863.873 | 7739.919 | 3.922588 | 0.020045687 |
| + |  | + | + |  |  |  |  | 6 | -3864.052 | 7740.278 | 4.281526 | 0.016752461 |
| + | + | + |  | + |  |  |  | 7 | -3863.027 | 7740.287 | 4.290123 | 0.016680610 |
|  | + | + |  |  |  | + |  | 6 | -3864.202 | 7740.578 | 4.581449 | 0.014419530 |
|  | + | + | + |  |  |  |  | 6 | -3864.238 | 7740.650 | 4.653134 | 0.013911856 |
| + | + |  | + |  |  |  |  | 6 | -3864.258 | 7740.690 | 4.693110 | 0.013636541 |
| + | + |  | + | + |  |  |  | 7 | -3863.366 | 7740.965 | 4.967889 | 0.011886027 |
| + | + | + |  |  | + | + |  | 8 | -3862.513 | 7741.324 | 5.327626 | 0.009929350 |
| + | + | + | + |  | + |  |  | 8 | -3862.513 | 7741.326 | 5.329505 | 0.009920029 |
| + | + | + |  |  |  | + |  | 7 | -3863.846 | 7741.924 | 5.927215 | 0.007357355 |
| + | + | + | + | + | + |  |  | 9 | -3861.778 | 7741.930 | 5.933268 | 0.007335122 |
| + | + | + |  | + | + | + |  | 9 | -3861.789 | 7741.952 | 5.955215 | 0.007255073 |
| + | + | + | + |  |  |  |  | 7 | -3863.873 | 7741.978 | 5.980959 | 0.007162282 |

**Table S11.** Model averaging table for the LMM for the peak frequencies of songs as the output of dredge function in R. isl: island, loc: territory location, trt: treatment, ord: treatment order. Plusses indicate that the term was included in that model. Only the models with delta AICc <6 are shown in the table. Intercept and bird ID as the random effect was included in each model. ICC for the random effect was 0.15 in the top model.

| isl | loc | trt | ord | Isl:loc | Isl:trt | loc:trt | Isl:loc:trt | df | logLik | AICc | delta | weight |
| --- | --- | --- | --- | --- | --- | --- | --- | --- | --- | --- | --- | --- |
| + |  | + | + |  | + |  |  | 7 | -3317.716 | 6649.663 | 0.000000 | 0.29388199 |
| + |  | + | + |  |  |  |  | 6 | -3319.535 | 6651.243 | 1.579652 | 0.13340001 |
| + | + | + | + |  | + |  |  | 8 | -3317.713 | 6651.725 | 2.061707 | 0.10482841 |
|  |  | + | + |  |  |  |  | 5 | -3321.000 | 6652.124 | 2.460655 | 0.08587141 |
| + | + | + | + |  | + | + |  | 9 | -3317.454 | 6653.283 | 3.619336 | 0.04811097 |
| + | + | + | + |  |  |  |  | 7 | -3319.535 | 6653.301 | 3.638068 | 0.04766246 |
| + | + | + | + | + | + |  |  | 9 | -3317.666 | 6653.706 | 4.043001 | 0.03892659 |
| + | + | + | + | + | + | + | + | 11 | -3315.587 | 6653.725 | 4.062120 | 0.03855625 |
|  | + | + | + |  |  |  |  | 6 | -3320.925 | 6654.025 | 4.361292 | 0.03319942 |
| + | + | + | + |  |  | + |  | 8 | -3318.924 | 6654.147 | 4.483475 | 0.03123192 |
|  | + | + | + |  |  | + |  | 7 | -3320.221 | 6654.674 | 5.010689 | 0.02399472 |
| + | + | + | + | + | + | + |  | 10 | -3317.418 | 6655.296 | 5.632656 | 0.01758155 |

**Table S12.** Model averaging table for the LMM for the duration of songs as the output of dredge function in R. isl: island, loc: territory location, trt: treatment, ord: treatment order. Plusses indicate that the term was included in that model. Only the models with delta AICc <6 are shown in the table. Intercept and bird ID as the random effect was included in each model. ICC for the random effect was 0.31 in the top model.

| isl | loc | trt | ord | Isl:loc | Isl:trt | loc:trt | Isl:loc:trt | df | logLik | AICc | delta | weight |
| --- | --- | --- | --- | --- | --- | --- | --- | --- | --- | --- | --- | --- |
| + |  | + |  |  | + |  |  | 6 | 2.665056 | 6.843801 | 0.000000 | 0.33149703 |
| + |  | + | + |  | + |  |  | 7 | 2.905835 | 8.420695 | 1.576894 | 0.15068204 |
| + | + | + |  |  | + |  |  | 7 | 2.684664 | 8.863037 | 2.019236 | 0.12078364 |
| + | + | + |  |  | + | + |  | 8 | 3.403725 | 9.491925 | 2.648125 | 0.08819555 |
| + | + | + |  | + | + |  |  | 8 | 2.974855 | 10.349666 | 3.505865 | 0.05743686 |
| + | + | + | + |  | + |  |  | 8 | 2.921396 | 10.456585 | 3.612784 | 0.05444695 |
| + | + | + |  | + | + | + |  | 9 | 3.742215 | 10.890569 | 4.046769 | 0.04382632 |
| + | + | + | + |  | + | + |  | 9 | 3.486631 | 11.401738 | 4.557937 | 0.03394189 |
| + | + | + |  | + | + | + | + | 10 | 4.308127 | 11.843037 | 4.999236 | 0.02722133 |
| + | + | + | + | + | + |  |  | 9 | 3.195476 | 11.984049 | 5.140248 | 0.02536816 |
| + | + | + | + | + | + | + |  | 10 | 3.810802 | 12.837687 | 5.993886 | 0.01655479 |
